# Regulations of expressions of rat/human sulfotransferases (SULTs) by anti-cancer drug, nolatrexed and micronutrients

**DOI:** 10.1101/2020.04.19.049007

**Authors:** Smarajit Maiti, Sangita MaitiDutta, Guangping Chen

**Author notes:** **Correspondence:** Dr. Smarajit Maiti, Professor and Head, Post Graduate Department of Biochemistry and Biotechnology, Cell & Molecular Therapeutics Lab, OIST, Midnapore-721102, E. Mail.

## Abstract

Cancer is a disease related to cellular proliferative-state. Drastically increase in cell-cycle regulations augments cellular folate-pool and folate-metabolism. So, this pathway is targeted therapeutically. A number of drugs are involved in this metabolism i.e. folic-acid/folinic-acid/nolatrexed(NT)/ methotrexate(MTX) for the research and treatment of cancer. Our previous study showed that MTX significantly modulated rat/human SULTs. Present study was an attempt to study the effect of NT (widely used in different cancers) and these micronutrients on the expressions of rat/human SULTs. Male Sprague-Dawley rats were treated with NT (01,10 or 100 mg/Kg) or both sexes were treated to folic acid (100,200 or 400 mg/kg) for 2-weeks and their AST-IV (2-napthol sulfation) and STa (DHEA-sulfation) activities, protein-expression (Western-Blot) and mRNA-expression (RT-PCR) were tested. In cultured HepG2 cells NT (1nM-1.2mM) or folonic-acid (10nM-10μM) were applied for 10 days. Folic acid (0-10μM) was treated to human hepatocarcinoma (HepG2) cells. PPST (phenol-catalyzing), MPST (dopamine) DHEAST (dehydroepiandrosterone,DHEA) and EST (estradiol-sulfating) protein-expressions (Western-blot) were tested in all HepG2 study. Present results suggest NT significantly increased SULTs expressions in rat (protein/mRNA/activity) and HepG2 cells. Folic acid increased SULTs activity/protein in sex-dependant manner (ASTIV in female/ STa in male). Both folic and folinic acid increased several hSULTs isoforms with varied level of significance (least or no increase at highest-dose) in HepG2 cells pointing its dose-dependent multi-phasic responses. The clinical importance of this study may be furthered in the verification of sulfation-metabolism of several exogenous/endogenous molecules, drug-drug interaction and their influences on the patho-physiological processes. Further studies are necessary in this regard.

## Introduction

Phase–II drug metabolising enzymes are important for their role in the modification of endogenous and exogenous compounds. Sulfotransferases are one group of enzymes catalyzes sulfation mediated polarization of drugs that increase solubility hence, the bioavailability of that compound (1–3). During this process biotransformation of any drug, bioactivation of that drug may occur. These steps may generate mutagenic or carcinogenic potentials in some drugs. From pro- to proximate and then to ultimate carcinogenic potentials of a drug may develop through its phase II metabolic pathway (4). The process of tumorigenesis and carcinogenesis follow few steps of impairments at cellular and /or metabolism of some endogenous molecule. The best example is estradiol (E_2_) (4, 5). Reports reveal that post menopausal women may develop tumor due to the higher level of E_2_ with low level of estrogen sulfotransferase (SULT1E1) (6, 7).

Low SULT1E1 fails to deactivate E_2_ to E_2_S (estrogen sulphate), so high E2 participate in cellular malfunctioned. Some bioamines or polyphenolic compounds are also associated with sulfation-desulfation related carcinogenic manifestation (6). Drugs used in cancer therapy are of great interest to study their potentials in the expression of SULTs. And whether the sulfation metabolism of drugs or its SULTs induction potentiality has some influence on cancer pathogenesis may be a subject of concern. Our laboratory and some other investigators report revealed that tamoxifen (TAM); an anti-estrogenic drug has SULTs regulation potentials. Further, therapeutic materials/ prescription drug mediated alterations of SULTs expression may also influence the sulfation of important endogenous molecules. A large number of prescription drugs are used in cancer therapeutics. And sometimes these drugs are applied for years and decades. Long term consumption of these drugs may have effects on sulfation metabolism of endogeneous molecules. So, studies on cancer drugs or vitamins or micronutrients (those have direct cell cycle regulatory potentials) are important. Our previous studies showed that cancer drug TAM and methotrexate (MTX) can alter SULTs expression (1,8). 4OH-TAM metabolism by sulfation has also been demonstrated. Some other report showed that cancer drug may alter SULTs expression. More or less related issues like apoptosis inducing drug Rhein and Emodin may alter SULTs expression (9).

In this background our present study is intended to elucidate the role of folic acid, folinic acid, nolatrexed on the expressions of human and rats SULTs. This study will help to reveal the role of prescription-drugs on SULTs expression. Moreover, alterations of SULTs expression how influence the sulfation metabolism of endogenous and exogenous drugs, and their physiological and pathological consequences and drug-drug interactions may also be explored.

## Materials and Methods

### Experimental Procedure

β-naphthol, [14C]b-naphthol (4.7 mCi/mmol), ρ-nitro-phenyl sulfate (PNPS), 3’-phosphoadenosine-5’-phosphosulfate (PAPS), and [1,2,6,7-3H(N)]dehydroepiandrosterone ([3H]DHEA, 60 Ci/ mmol) were purchased from Sigma-Aldrich (St. Louis,MO). SDS–polyacrylamide gel electrophoresis reagents were obtained from Bio-Rad (Hercules, CA). Western blot chemiluminescence reagent kits (Super Signal West Pico Stable Peroxide and Super Signal West Pico Luminol/ Enhancer solutions) were purchased from Pierce Chemical (Rockford, IL). Nitrocellulose membrane (Immobilon-P; Millipore Corporation, Bedford, MA) used during Western blot procedure was purchased from Fisher Scientific (Fair Lawn, NJ). Total RNA extraction kit (RNeasy mini protection kit) was supplied by Qiagen (Valenica, CA). Antibodies against ASTIV (12) and STa (11) were provided by Dr. Michael W. Duffel (Division of Medicinal and Natural Products Chemistry, College of Pharmacy, The University of Iowa, Iowa City, IA). Protein assay reagent was purchased from Bio-Rad. All other reagents and chemicals were of the highest analytical grade available.

### Animals and drug treatment

Female Sprague–Dawley rats (Harlan, Indianapolis, IN) 10- to 11-weeks old and 200–300 g body weight were used in this investigation. Rats were housed in a temperature and humidity-controlled room and supplied with rodent chow and water for at least 1 week before use. Rats of both sexes were divided into 4 groups with 3 in each. NT was suspended in corn oil and administered by gavages at 1, 10,100 mg/kg/day for 2 weeks to 3 separate groups of female rats. Folic acid was and administered by gavages at (100, 200 or 400 mg)/kg/day for 2 weeks to 3 separate groups of both male and female rats. The corresponding group of control rats received only the vehicle. The animals were sacrificed 24 hours after the final drug treatment. Livers were collected, washed with sterile, ice-cold NaCl (0.9%, w/v) solution, and kept in dry ice bath. Samples were stored at-80°C until use.

### Cell culture and drug treatment

HepG2 cells were obtained from the American Type Culture Collection (Manassas, VA). HepG2 Cells were grown and maintained in Dulbecco’s Modified Eagles’s Medium Nutrient Mixture F-12 Ham (Sigma) supplemented with L-glutamine and 15mMHEPES, and 10% fetal bovine serum (FBS). The cultures were incubated at 37°C in a humidified incubator containing 5% CO2, 95% air (9). After seeding at 0 days, on day 1, FA (0.01, 0.1, 1, 10 μM final) was added to the medium in properly marked plates. Similarly, for the NT experiment, NT was added (1ηM to 1.2 mM) to the properly marked plates carrying HepG2 cells. Control plates are added with the vehicle. The medium was refreshed every 3 days with the new addition of corresponding drug. On day ten the cells were harvested. Cytosols are prepared from the cells by suitable methods as explained (9).

### Cytosolic sample preparation

Liver homogenates were prepared with 50mM Tris buffer containing 0.25M sucrose, pH 7.5. Homogenates were centrifuged at 100,000g for 1 h at 4°C Cytosol aliquots were collected and preserved at 80 °C for enzymatic assay and Western blot.

### Sulfotransferases (SULTs) assay

Two different enzyme assay methods were used.

### PNPS assay method

β-Naphthol sulfation activity from liver cytosols was determined as previously described (10,11,12). This assay determines phenol sulfation activities of different isoforms of phenol sulfating SULTs. Briefly, sulfation activity was determined in a reaction mixture containing 50mM Tris buffer, pH 6.2, 5mM PNPS, 20 lM PAPS, and 0.1mM β-naphthol. Rat liver cytosols (50 μg protein) were used as the enzyme source in a total reaction volume of 250 μl. After 30 min incubation at 37°C in a shaking water bath, the reaction was stopped by adding 250μl of 0.25M Tris, pH 8.7. The reaction mixtures were read at 401 nm in a spectrophotometer. Specific activity (SA) was expressed as nanomoles per minute per milligram of protein. The data shown in the figures are the average of 3 independent data sets collected from 5 different animals.

### Radioactive assay method

β-Naphthol sulfation activity in intestinal cytosols and DHEA sulfation activities in liver cytosol was determined by the radioactive assay method previously described (12). Other ingredients and reaction conditions were same as the PNPS assay mentioned above. For intestinal β-naphthol sulfation activity, [14C]β-naphthol (4.7 mCi/mmol; 0.1mM final concentration) was used as substrate. To determine DHEA sulfation activity in liver cytosol, [3H] DHEA (diluted to 0.4 Cμ/mmol; 2 μM final concentration) was used as substrate. For all assays, 20 μM PAPS was used. Liver cytosol protein (50 μg) was used as enzyme source in a total reaction volume of 250 μl. After 30 min incubation at 37 °C in a shaking water bath, the reaction was stopped by adding 250 μl of 0.25M Tris, pH 8.7. Extraction was performed twice by addition of 0.5 ml of water-saturated chloroform. After the final extraction, 100 μl of aqueous phase was used for scintillation counting. The data shown in the figures are the average of 3 independent data sets collected from 3 different animals. PAPS was eliminated from the controls of both assay methods. Assays were run in duplicate and the average of the results was used for enzyme activity calculations.

### Western blot analysis for the detection of SULTs

Cytosol protein from liver (10 μg) was used in a 10% polyacrylamide gel in an electrophoresis system (Novex, San Diego, CA). After running at 200 V, the protein bands were transferred overnight at 40V onto a nitrocellulose membrane. For rat liver cytosols, membranes were incubated with either rabbit anti-rat AST-IV or rabbit anti-rat STa (1:5000) in TBST (50mM Tris, pH 7.5, 150mM NaCl, and 0.05% (v/v) Tween 20) containing 5% (w/v) dried milk for 2 h on a shaker at room temperature. For Hep G2 cell cytosols, membranes were incubated with rabbit anti-hPPST or hMPST or hDHEAST or hEST (1:5000 to 1:2000) antibodies. After incubation, all membranes were washed with TBST for 4-15 min and incubated in secondary antibody (horseradish peroxidase-conjugated Immuno-Pure goat anti-rabbit IgG; H+ L) at 1:5000 dilutions in the same buffer for 2 h. The membranes were washed with TBST for 4-15 min and then with phosphate-buffered saline (PBS) 3-5 min. Fluorescent bands were developed with 1ml of substrate containing the same volume of each Super Signal West Pico Luminol Enhancer solution and Super Signal West Pico Stable Peroxidase solution at room temperature for 5 min. The X-ray films were exposed to the membrane and then developed. Films were scanned and the densitometric analysis was performed in a Gel Documentation and Analysis System from Advanced American Biotechnology and with AAB software (Fullerton, CA).

### Extraction of total RNA and RT-PCR for SULTs mRNA expression study

Total RNA was extracted from liver using RNeasy mini protection kit from Qiagen according to supplier’s guidelines. The concentration and purity of the extracted RNA were checked spectrophotometrically by measuring 260/280 absorption ratios. The primer pair for AST-IV was designed in our laboratory using the Gene Fisher primer designing and Multialignment software. Using the forward primer (FP) 50-GTGTCCTATGGGTCGTGGTA-30 and reverse primer (RP) 50-TTCTGGGCTACAGTGAAGGTA-30 (GenBank Accession No.: X52883), the 299-bp AST-IV cDNA was synthesized (10). The 264-bp STa cDNA was synthesized using the primer pair FP 50-TCCTCAAAGGATATGTTCCG-30 and RP 50-CAGTTCCTTCTCCATGAGAT-30 (GenBank Accession No.: M33329) (10). The nucleotide sequences X52883 and M33329 (GenBank Accession No.) were used as reference sequence for the synthesis of AST-IV and STacDNA, respectively. The specificity of all primers was tested using the BLAST of the National Center for Biotechnology Information Open Reading Frame software. cDNA synthesis from 1 μg of liver total RNA was performed in a 50 μl reaction mixture. The concentrations of the different ingredients were used following supplier’s protocol. For control, 500-bp cDNA of rat b-actin was synthesized from the same amount of RNA from respective tissues. The primer pair (FP 50-GATGTACGTAGCCATCCA-30 and RP 50-GTGCCAACCAGACAGCA-30) for the synthesis of rat *β*-actin cDNA was designed in our laboratory using the same software mentioned above (10).

### Statistical analysis

Student_s t test was performed to calculate the statistical significance of the difference s between means of control and drug-treated rats or cultured cells. Data presented in the figures denote means±SEM of the results collected separately from five individual experiments.

## Results and Discussion

Present results suggests that NT significantly increased 2-napthol sulfation (SULT1A1 or ASTIV, p<0.01 and p<0.05) and DHEA sulfation (SULT2A1 or STa, p<0.01 and p<0.05) activities in rat liver with all drug doses except the highest dose in case of SULT2A1 activity (fig 1). NT is widely used as an anti-cancer drug in last decade against a number of cancers. Here, we first time report the NT induced SULTs expressions and altered their activities in human and rat cells. This work has notable importance in relation to pathological/ pharmacological sciences. Anti-metabolites are active chemotherapeutic agents for many solid tumor and hematologic malignancies (13). Folate antagonists, purine analogues, and pyrimidine analogues are the three main categories of anti-metabolites (14). Methotrexate, the most studied folate antagonist (inhibitor of dihydrofolate reductase, DHFR), is effective in many malignancies (15). However, resistance to MTX develops by decreased folate carrier-mediated membrane transport (1). Our previous study suggested that MTx can induce different SULTs in rat liver. Supplementation with folic acid and vitamin B12 has been shown to reduce the toxicity of pemetrexed without affecting efficacy and has increased the therapeutic index for this novel agent (13). This may indicate MTx related sulfation metabolism and unwanted events may be counteracted by the folate supplementation, when it is more efficient in the presence of B12 for nucleotide metabolism.

**Fig. 1.**
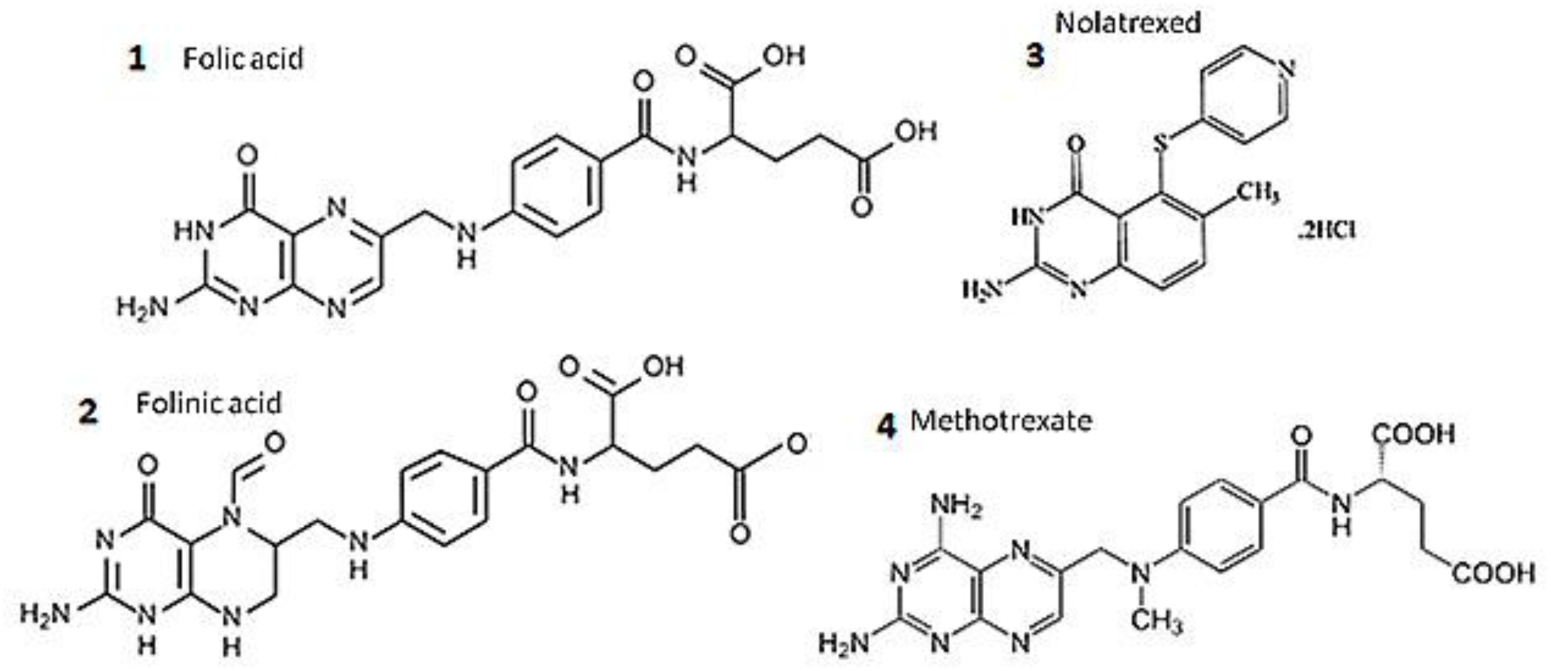
Chemical structures of anticancer drugs and micronutrient, folic acid

Regulation of expressions of phase-II enzymes especially, SULTs by prescription drugs are important to learn the influences of different sulfation reactions in patho-physiological conditions. Nolatrexed dihydrochloride, a thymidylate synthase inhibitor was tested to establish the most tolerable dose for Phase II studies in a number of patients and found to be safe up to a certain dose (16), but its role on sulfation metabolism has not been verified earlier. In the current study, it has been shown that NT may induce rat and human SULTs isoforms at protein and RNA level (fig 2). Two randomized trials (one in the USA and other in Europe) demonstrated a brief comparison between methotrexate (MTX) and nolatrexed (NT) in patients with certain type of cancer. This study demonstrated similar pattern of response, prognosis and overall survival period for both the drugs (17). In our previous studies we have clearly demonstrated changes in SULTs expressions by MTx in rat tissues and in human cultured cells (11). In some trial, nolatrexed was administered with as high as the dose of 725 mg/m (2)/day to hepatocellular carcinoma patients without any toxicity symptom. Our present experimental dose to rat or human HepG2 cell is far lower than the clinical dose. Neither with our dose (data not shown) nor with the clinical doses any cumulative toxicity was noticed (18). No systemic investigation has been shown NT effect on SULTs expression. Both the drugs NT and MTX have been shown to be associated to folic acid and folinic acid with the structural/functional analogy (fig 1).

**Fig-2.**
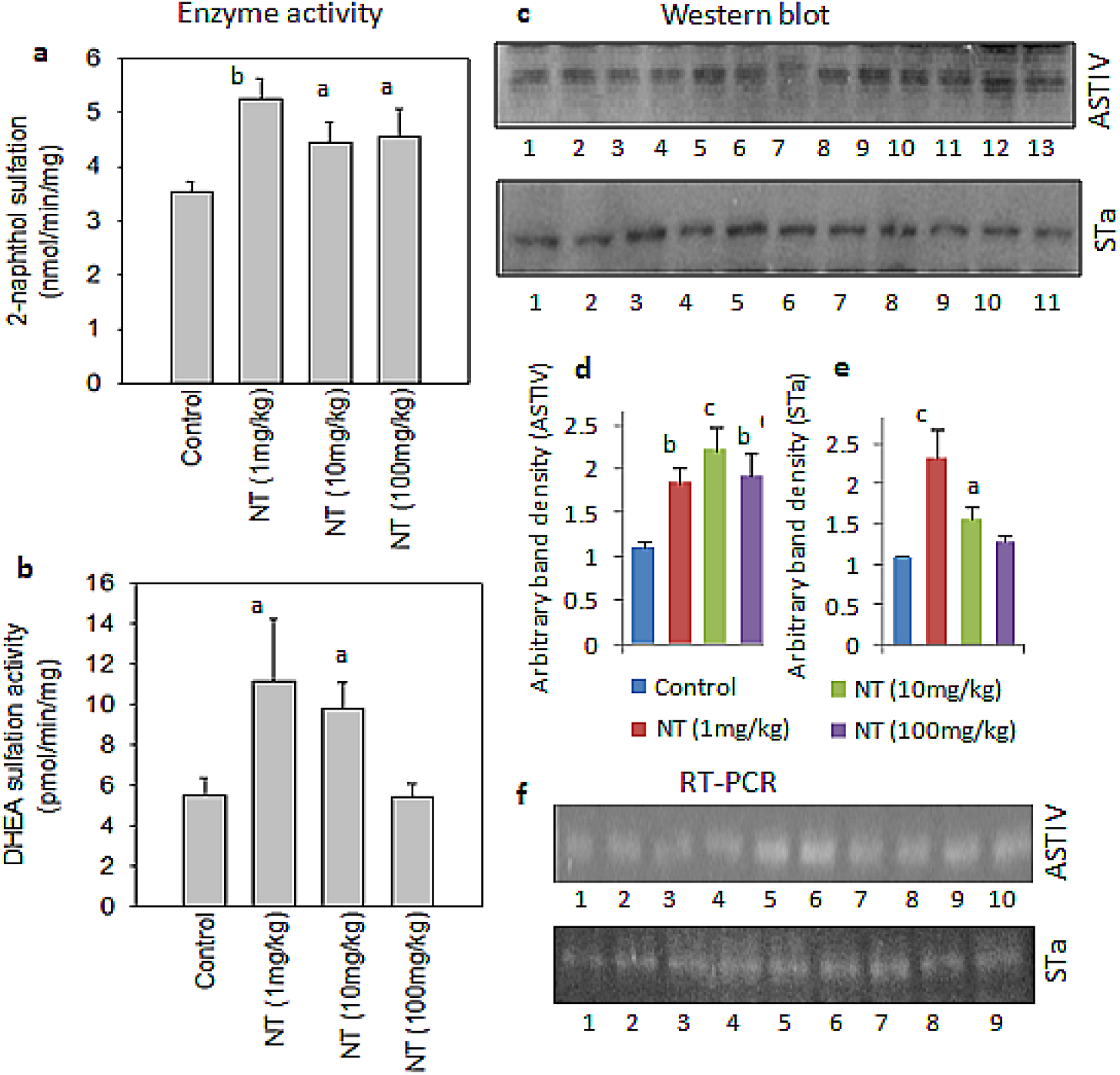
Studies of female rat experimental model demonstrate ASTIV and STa activity and in liver tissue increased by the NT treatment for 2 weeks (a,b). AST IV and STa protein expressions results from Western blot analysis are presented (c) and their densitometry analysis are drawn as bar-diagram (d,e). Western blot study on this protein expression and their densitometry analysis study basically agree with the enzymatic activity data. Lane distribution: AST IV: 1-3 = control, 4-6 =NT 1mg/Kg, 7-9 = 10mg/Kg, 10-13 = 100mg/Kg. STa: 1-2 = control, 3-5 =NT 1mg/Kg, 6-8 = 10mg/Kg, 9-11 = 100mg/Kg. RT-PCR data of ASTIV and STa are presented in figure 2e. Though there are some inconsistencies this basically supports the Western blot results. Lane distribution: AST IV: 1-3 = control, 4-6 =NT 1mg/Kg, 7,8 = 10mg/Kg, 9,10 = 100mg/Kg. STa: 1-2 = control, 3-5 =NT 1mg/Kg, 6,7 = 10mg/Kg, 8,9 = 100mg/Kg. Results in bar diagram represent mean±SE of 5 independent sample /experiment. Level of significance is represented ^a^ P<0.05, ^b^ P<0.01, ^c^P <0.001

Xenobiotic induction of SULTs is not well known. Our enzyme assay, Western blot, and reverse transcription polymerase chain reaction (RT-PCR) results demonstrated that protein and mRNA expressions of aryl sulfotransferase (AST-IV) and hydroxysteroid sulfotransferase (STa) were induced in liver and intestine of male and female rats following MTX (1). Our earlier investigation showed that MTX is an inducer of rat and human sulfotransferases (1). Further, we report that folic acid treatment inhibited MTX induction of aryl sulfotransferase (AST-IV) in female rat liver and hydroxysteroid sulfotransferase (STa) in male rat liver. This is important for understanding the clinical mechanisms of MTX (12). In the current study, in addition to the SULT induction in rat by NT, this drug inconsistently induced DHEAST in human HepG2 cell. But with the same treatment schedule EST expression was moderately inhibited. At a low (ηM) to high (mM) dose multiphasic action of NT is noticed (fig 3). This should be taken into account at the time of clinical use of this drug for a longer period. Sex dimorphic increase of SULTs expressions are noticed in rat liver in response to folic acid treatment. The basal level of ASTIV and STa are found to be lower in female and male rats, respectively (fig. 2), but the drug induced increase in these enzymes are found to be in reverse order (p<05 to p<0.001). These findings are supported by some earlier reports (1,8). Folic acid was noticed to be inducing the PPST, MPST and EST expressions in human carcinoma HepG2 cells. This suggests that FA or its analog or related drugs have similar pattern on influence on SULTs expressions and activities. The overall changes on SULTs activities might have some influence on the sulfation metabolism of several biomolecules and endo- or xenobiotics. Continuation of this study is required in this regard. Not only for adult cancers, but NTs use for children cancer is reported to be safe. So the therapeutic activity is correlated to its antiproliferative toxicity (19). For metastatic hepatocellular carcinoma (HCC), treated with nolatrexed (NT) or doxorubicin (DOX) showed minimal activity in this phase III trial (20). The pathophysiological outcome of NT in relation to different cancer has not been enlightened with its role on phase I and phase II drug metabolizing enzymes. We want to draw the attention that what is explained as ‘safe use’ may be extrapolated as the most effective and optimized use of this drug. And for this sulfation metabolism study by NT is important.

**Fig-3.**
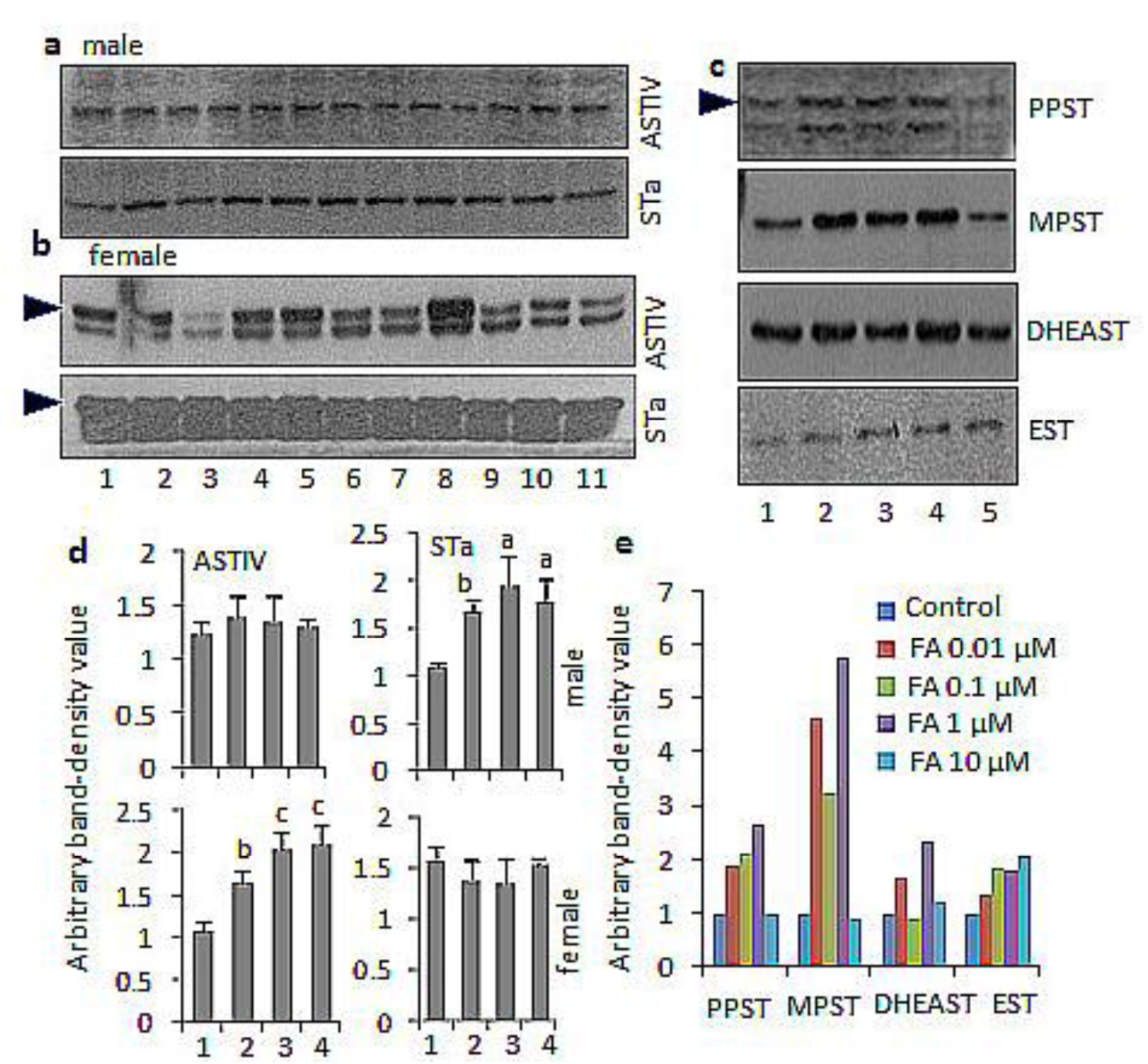
Sex dimorphic protein expression (ASTIV and STa) were noticed after Folic acid exposure to experimental rat. FA –dose dependent increase in STa in male and ASTIV in Female rat liver were noticed (a). Treatment of FA to Hep-G2 cells suggests consistent increases of PPST, MPST and gradual increase in EST expression. At highest 10μM of FA-dose PPST, MPST and DHEAST did not respond significantly (b). Results in bar diagram represents densiometry data (d) ASTIV and STa, bar diagrams represent mean±SE of 5 independent experiments. Level of significance is represented ^a^P<0.05, ^b^P<0.01, ^c^P<0.001

**Fig. 4.**
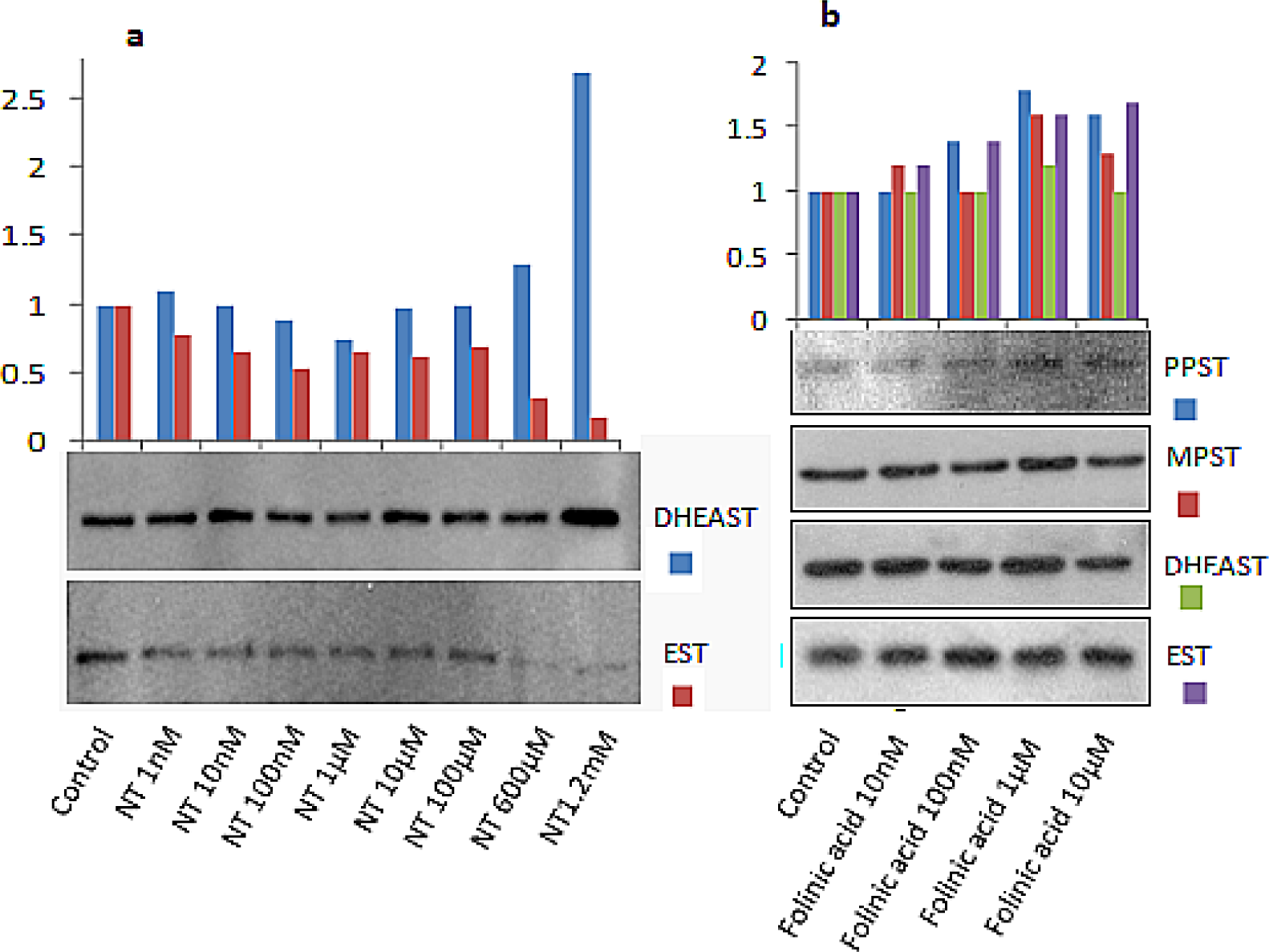
Dose responsive increases of DHEAST in HepG2 cells to very low level of NT. At highest 1.2 mM NT concentration, two and half fold increase of DHEAST was noticed. Gradual decrease of EST was noticed in these cells after same dose of NT application (a). Folinic acid increased PPST, DHEAST and EST in dose responsive manner. The fold of increase was found to be less at the highest concentration of folinic acid.

With reference to our previous discussion, it is notable that cancer chemotherapy is a combination of multiple drug regimens where drug-drug interaction is possible. And this interaction may influence the therapeutic mechanism of that drug. As for example, cantharidin and nolatrexed were used to inhibit PP and TS activity, respectively. Synergistic manner of response of these drugs are more effective than only NT use (21). Further studies are necessary to explore whether this effect or possible toxicities are imposed by drug-induced sulfation-metabolism or not. Similarly pharmacokinetics of nolatrexed is more therapeutically favorable when it is administered sequentially with paclitaxel (22). These schedules of uses necessitate the study of its impact on SULTs expressions to evaluate the possible sulafation metabolism of other drugs and further possible events of drug-drug interactions. Metabolic reprogramming of tumor cells toward serine catabolism is now recognized as a hallmark of cancer (23). These observations provide insights into the mechanism of action of antifolate drugs and that may help to more rational drug-designing (24).

Drug resistance is the main barrier to more effective treatment of cancers with antifolates; therefore, mechanisms of antifolate resistance are the topic of worth for investigation (25). As because, thymidylate synthase (TS) is a nucleotide and cell cycle regulator so TS-targeted chemotherapy may impact on pharmacogenetics which is considered in selection of 5-Flurouracil therapy for colorectal cancer (26). Similarly pharmacogenomic influence by NT also should be verified. Both folate and methotrexate transport were inhibited by classical antifolates but not by non-classical antifolates or biopterin (27). Not only the drug-metabolizing enzyme genes but also the role of tumor suppressor genes (i.e. p53) has been important to alter the sensitivity to TS inhibitors which accounts for the outcome of chemotherapy (28). In further search on the mechanism of MTX inductions of SULTs expression our previous studies explained in involvement of several nuclear receptors like CAR and PXR (29,30). And these and other report justifies molecular impact of antifolate drugs is occurring at the transcriptional signalling level.

In conclusion, here for the first time we demonstrate the role of NT on SULTs expression and induction of their increased activities. In our earlier studies SULTs inductions by methotrexate and tamoxifen have been shown. All are anti-cancer drug. This induction may have some unidentified impact on sulfation metabolism of different molecules, drug-drug interaction with physiological and pathological consequences. Further studies are necessary in this regard.

## Acknowledgements

Institutional members

## Notes

**Conflict of interests:** None

### Competing Interest Statement

The authors have declared no competing interest.

